# Evolutionary conservation and multilevel post-translational control of S-adenosyl-homocysteine-Hydrolase in land plants

**DOI:** 10.1101/2019.12.20.884296

**Authors:** Sara Alegre, Jesús Pascual, Andrea Trotta, Martina Angeleri, Moona Rahikainen, Mikael Brosche, Barbara Moffatt, Saijaliisa Kangasjärvi

**Affiliations:** Department of Biochemistry, Molecular Plant Biology, University of Turku, FI-20014 Turku, Finland; Organismal and Evolutionary Biology Research Program, Faculty of Biological and Environmental Sciences, Viikki Plant Science Centre, University of Helsinki, FI-00014 Helsinki, Finland; Department of Biology, 200 University Avenue, University of Waterloo, Ontario, Canada, N2L 3G1

## Abstract

Trans-methylation reactions are intrinsic to cellular metabolism in all living organisms. In land plants, a range of substrate-specific methyltransferases catalyze the methylation of DNA, RNA, proteins, cell wall components and numerous species-specific metabolites, thereby providing means for growth and acclimation in various terrestrial habitats. Trans-methylation reactions consume vast amounts of S-adenosyl-L-methionine (SAM) as a methyl donor in several cellular compartments. The inhibitory reaction by-product, S-adenosyl-L-homocysteine (SAH), is continuously removed by SAH hydrolase (SAHH) activity, and in doing so essentially maintains trans-methylation reactions in all living cells. Here we report on the evolutionary conservation and multilevel post-translational control of SAHH in land plants. We find that SAHH forms oligomeric protein complexes in phylogenetically divergent land plants, and provide evidence that the predominant enzyme is a tetramer. By analyzing light-stress-induced adjustments occurring on SAHH in *Arabidopsis thaliana* and *Physcomitrella patens*, we demonstrate that both angiosperms and bryophytes undergo regulatory adjustments in the levels of protein complex formation and post-translational modification of this metabolically central enzyme. Collectively, these data suggest that plant adaptation to terrestrial environments involved evolution of regulatory mechanisms that adjust the trans-methylation machinery in response to environmental cues.

## INTRODUCTION

Land plants have evolved sophisticated biochemical machineries that support cell metabolism, growth and acclimation in various terrestrial habitats. One of the most common biochemical modification occurring on biological molecules is methylation, which is typical for DNA, RNA, proteins, and a vast range of metabolites. Trans-methylation reactions are therefore important in a relevant number of metabolic and regulatory interactions, which determine physiological processes during the entire life cycle of plants. Trans-methylation reactions are carried out by methyl transferases (MTs), which can be classified into O-MTs, N-MTs, C-MTs and S-MTs based on the atom that hosts the methyl moiety [1,2]. All these enzymes require *S*-adenosyl-L-methionine (SAM) as a methyl donor [3]. Among MTs, O-MTs form a large group of substrate-specific enzymes capable of methylating RNA, proteins, pectin, monolignols as well as various small molecules in various cellular compartments [2].

The availability of SAM is a prerequisite for methylation, while the methylation reaction by-product, S-adenosyl-L-homocysteine (SAH) is a potent inhibitor of MT activity and must therefore be efficiently removed [4]. To ensure the maintenance of SAM-dependent trans-methylation capacity, SAH is rapidly hydrolysed by *S*-adenosyl-L-homocysteine hydrolase (SAHH, EC 3.3.1.1) in a reaction that yields L-homocysteine (HCY) and adenosine (ADO) [5]. Subsequently, methionine is regenerated from HCY by cobalamin-independent methionine synthase (CIMS, EC 2.1.1.14) using methyltetrahydrofolate as a methyl donor. Finally, methionine is converted to SAM in an ATP-dependent reaction driven by SAM synthase, also known as methionine adenosyltransferase (SAMS/MAT, EC 2.5.1.6). Together these reactions are termed the Activated Methyl Cycle (AMC).

SAHH is the only eukaryotic enzyme capable of hydrolysing SAH, and therefore it is a key player in the maintenance of cellular transmethylation potential. The *Arabidopsis thaliana* (hereafter Arabidopsis) genome encodes two SAHH isoforms, SAHH1 (AT4G13940) and SAHH2 (AT3G23810); SAHH1 is indispensable for physiological functions at different developmental stages [6–8]. Null mutations of SAHH1 are embryo lethal in Arabidopsis and severe symptoms caused by SAHH deficiency have been reported in human [6,9,10] whereas there were no morphological abnormalities in homozygous Arabidopsis *sahh2* T-DNA insertion mutants [6]. Mutants suffering from impaired SAHH1 function, including the knock-down *sahh1* and *homology-induced gene silencing 1* (*hog1*), are viable but have delayed germination, slow growth and short primary roots [6,11]. Evidently, SAHH is crucial in ensuring accurate metabolic and regulatory reactions in the cell. Even though a number of post-translational modifications (PTMs) has been detected on Arabidopsis SAHH [2], the exact regulatory impact of these modifications remain poorly understood.

At the amino acid sequence level, SAHH is one of the most highly conserved enzymes across the kingdoms of life [12]. Crystallography and structural studies from phylogenetically distant species have reported SAHH to form dimers, tetramers and hexamers in plant, mammalian and bacterial species [13–16]. The high-resolution crystal structure of *Lupinus luteus* SAHH1 suggested that in higher plants the enzyme would form functional dimers with a calculated molecular mass of 110 kDa [14,15]. However, in Arabidopsis leaves SAHH is predominantly present in an oligomeric protein complex called SAHH complex 4, which has an approximate molecular weight of 200 kDa [17]. The subunit composition of this abundant oligomeric SAHH complex has not been resolved.

Here we report on the evolutionary conservation and biochemical characteristics of SAHH in land plants. We find that the predominant oligomeric SAHH complex 4 can be detected in phylogenetically divergent land plants, and provide evidence suggesting that the protein complex is a tetrameric form of the enzyme. By analyzing regulatory adjustments occurring on SAHH in high-light-exposed Arabidopsis and *Physcomitrella patens* (hereafter Physcomitrella), we demonstrate that both angiosperms and bryophytes respond to light-induced stress by regulatory adjustments in this metabolically central enzyme.

## MATERIALS AND METHODS

### Plant material

*Arabidopsis thaliana* wild type ecotype Columbia-0, a transgenic *Arabidopsis* line stably expressing SAHH1p::EGFP-SAHH1 [18], *Brassica oleracea* convar. *acephala* varieties Half Tall and Black Magic (kales) and *Lupinus luteus* were grown in peat:vermiculite (2:1) and 50% relative humidity at 8-hour light period under 130 µmol photons m^-2^ s^-1^ and 22°C. Samples were collected after 4 weeks of growth. *Spinacea oleracea* (spinach) and *Brassica oleracea* convar. *italica* (broccoli) were purchased from the local supermarket. *Physcomitrella patens* was grown for 13 days on agar plates in minimum media [19] in a 16-hour photoperiod under 45 µmol photons m^-2^ sec^-1^ at 24°C. For high light stress experiments, Arabidopsis was grown for 16 days in a 12-hour light period under 130 µmol photons m^-2^ s^-1^ and thereafter shifted to 800 µmol photons m^-2^ s^-1^ at 26°C at a 12-hour light period for 2 days. *Physcomitrella* was grown as described above and shifter to 500 µmol photons m^-2^ s^-1^ in a 16-hour light period for 2 days.

### Analysis of publicly available transcript profiles

O-methyltransferases were selected from the UniProt database (https://www.uniprot.org/) using the search criteria: “O-methyltransferases” + “*Arabidopsis thaliana”* + “reviewed” (September 2019). This list was supplemented with Activated Methyl Cycle enzymes according to Rahikainen et al. (2018). Together, the selected O-MTs and AMC enzymes formed a total of 40 genes, which were used as the input in GENEVESTIGATOR [20]. This input was however assigned to 39 genes since AT5G17920 and AT3G03780, encoding CIMS1 and CIMS2, respectively could not be distinguished because they share the same probe in Affymetrix Arabidopsis ATH1 microarray. The database search was limited to “Only Columbia-0 Wild Type from Affymetrix Arabidopsis ATH1 genome array”. The “perturbations” tool from GENEVESTIGATOR was used to determine in which experimental conditions the selected genes were differentially expressed. Experiments in which at least 60% (24 out of 39) of the input genes were differentially expressed (p-value <0.05) were selected as data to generate the heatmap. The perturbations were hierarchically clustered with R package pheatmap (v1.0.12) [21] using Ward’s method and Euclidean distance.

### Confocal microscopy

Fluorescence from eGFP was imaged with a confocal laser scanning microscope Zeiss LSM780 with either C-Apochromat 40x/1.20 W Korr M27 or Plan-Apochromat 20x/0.8 objective. eGFP was excited at 488 nm and detected at 493 to 598 nm wave length and chlorophyll fluorescence was excited at 633 nm and detected at 647 to 721 nm wave length. Hectian strands were visualized by plasmolysis with 1 M NaCl and imaged after 6 minutes incubation. Images were created with Zeiss Zen 2.1 software version 11.0.0.190.

### Isolation of protein extracts and biochemical analysis of SAHH

Leaves of *A. thaliana, B. oleracea* convar. *acephala* (Half Tall and Black Magic), *B. oleracea convar. italica, L. luteus* and *S. oleracea*, and aerial parts of *P. patens* were ground in liquid nitrogen and mixed with extraction buffer [10 mM HEPES-KOH pH 7.5, 10 mM MgCl_2_, supplemented with protease (Pierce EDTA-free Minitablets; Thermo Fisher Scientific) and phosphatase (PhosSTOP; Roche) inhibitors]. The samples were centrifuged at 18 000 g for 15 minutes and the soluble fractions were taken for further analysis.

For biochemical analysis of Arabidopsis SAHH complex 4, soluble protein fractions were treated with 0.25 %, 1% sodium dodecyl sulfate (SDS; CAS Number 151-21-3) and/or 10 mM dithiothreitol (DTT; CAS Number 3483-12-3) in a total volume of 20 µL for 60 minutes as indicated in the figure legends. To assess protein complex formation, soluble protein fractions corresponding to 5 µg of protein were separated on Clear Native (CN) PAGE with a 7.5-12% gradient of acrylamide as in Rahikainen et al. (2017). Isoelectric focusing (IEF) was performed as in Lehtimäki et al. (2014). Phos-tag gel electrophoresis with 7.5% (w/v) acrylamide in the separation gel was performed according to manufacturer’s instructions (www.wako-chem.co.jp; https://labchem-wako.fujifilm.com/us/category/docs/00899_doc01.pdf). SAHH was detected by immunoblotting with anti-SAHH antibody or by using SYPRO as a protein stain as described in Trotta et al. (2011).

### Amino acid alignment and construction of phylogeny tree

SAHH amino acid sequences from *Arabidopsis thaliana* (accession AT4G13940; The Arabidopsis Information Resource, www.arabidopsis.org), *Lupinus luteus* (accession Q9SP37; https://www.uniprot.org), *Brassica oleracea* convar. *capitata*, (accession Bol033424; https://phytozome.jgi.doe.gov/pz/portal.html), *Spinacia oleracea* (accession A0A0K9RFV6; https://www.uniprot.org) and *Physcomitrella patens* (accession Pp3c19_13810V3.1; https://phytozome-next.jgi.doe.gov/) were aligned using ClustalW Multiple Alignment [24] in BioEdit Version 7.2.5. *Brassica oleracea* convar. *acephala* amino acid sequence was unavailable, thus *Brassica oleracea* convar. *capitata* was used as it is closest relative with available sequence. Identities and similarities were reckoned in BioEdit by pairwise comparison with BLOSUM62 matrix. A phylogenetic tree was built with the same amino acid sequences plus human SAHH1 (accession P23526; https://www.uniprot.org) using Neighbor-Joining method [25] in MEGA7.

## RESULTS

### Dynamics of O-MT mRNA abundance in *Arabidopsis*

The accuracy and biochemical specificity of trans-methylation reactions stem from a high number of substrate-specific MTs, whose expression patterns can be highly responsive to both endogenous and exogenous signals. Here we focused on AMC enzymes and the O-MTs, which are well known for their functions in the methylation of small metabolites that accumulate upon environmental perturbations. To illustrate the dynamism of O-MT transcript abundance in Arabidopsis, we performed an exploratory analysis using the “Perturbations” tool in GENEVESTIGATOR. Reviewed O-MTs from *A. thaliana* gene list were obtained from UniProt (www.uniprot.org; September 2019) and this list was supplemented with enzymes of the AMC (S1 Table). Perturbations in which at least 60% (24 out of 39) of the genes for the AMC enzymes and selected MTs were differentially expressed with a p-value <0.05 were selected to build the cluster heatmap.

As shown in Fig 1, hierarchical clustering of the gene expression data suggested dynamic adjustments in O-MT transcript abundance in response to a number of perturbations, which formed seven clusters. Clusters 6 and 7, branched out from the rest of clusters and included processes related to hormonal signalling and seed germination, respectively, while clusters 1 to 5 were comprised of perturbation categories related to light conditions, biotic, abiotic and chemical stress, and other physiological processes. These findings supported the view that O-MTs are highly regulated at the level of mRNA abundance and contribute to a multitude of metabolic processes in different compartments of plant cells.

**Fig 1.**
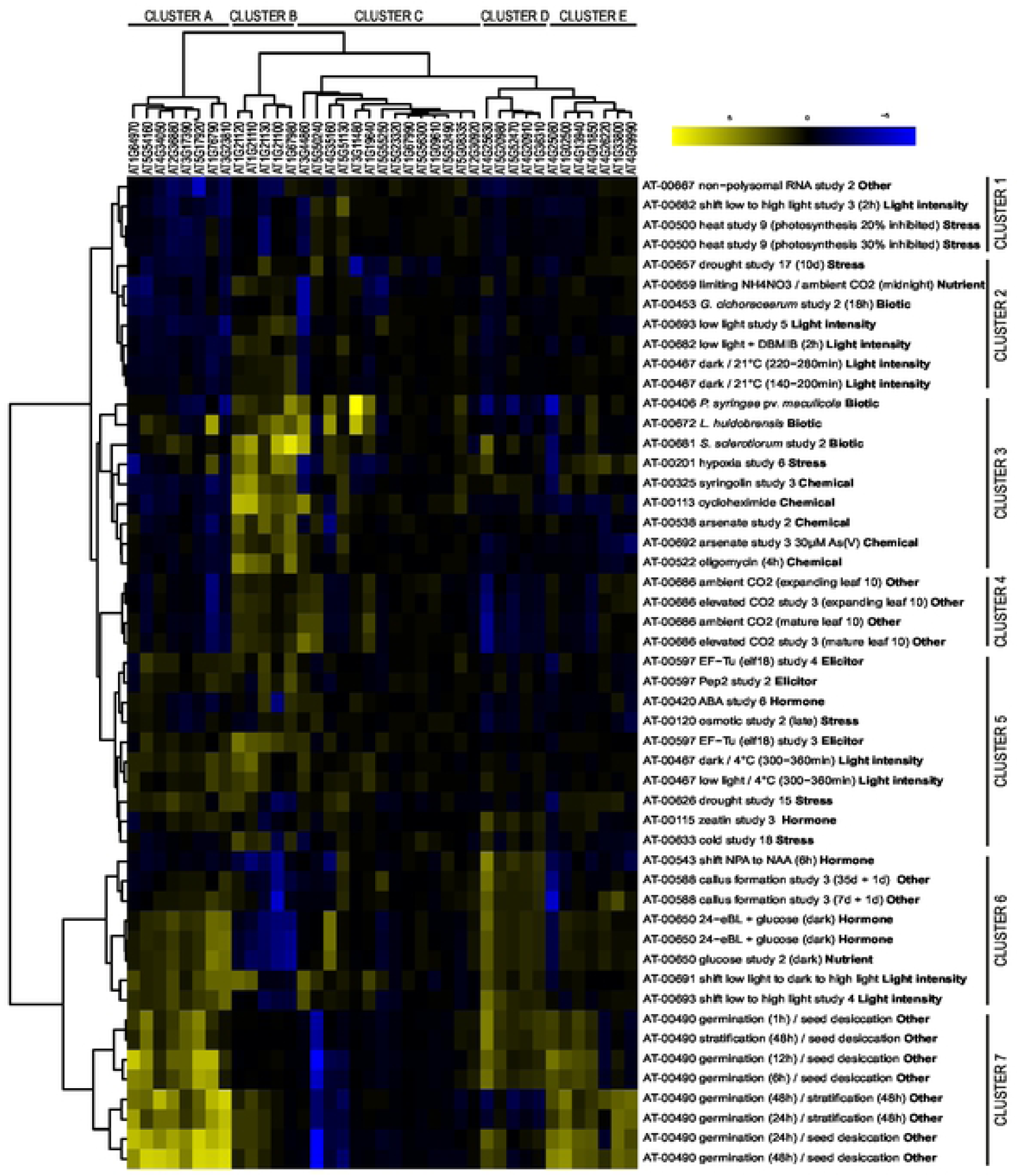
Hierarchically clustered heatmap depicting dynamic adjustments in the transcript abundance for genes encoding O-METHYLTRANSFERASES (O-MTs) and enzymes of the activated methyl cycle in Arabidopsis. The analysis was performed using the “Perturbations” tool in GENEVESTIGATOR. Experiments in which at least 60% of the input genes were differently expressed were selected to build the cluster heatmap. The input genes were retrieved from UniProt reviewed database by searching for “O-methyltransferase” and filtering for species (*Arabidopsis thaliana*), and this list was combined with genes encoding the AMC enzymes (S1 Table). *This gene represents both AT5G17920 and AT3G03780 as they were indistinguishable because they share the same probe in Affymetrix Arabidopsis ATH1 microarray chip.

### Sub-cellular localization of SAHH1 in Arabidopsis leaves

Next we assessed the sub-cellular localization of Arabidopsis SAHH1. Confocal microscopy imaging of leaves of four-week-old Arabidopsis plants stably expressing an EGFP-SAHH1 fusion protein under the native SAHH1 promoter (SAHH1p::EGFP-SAHH1) [18], revealed that SAHH localized to multiple sub-cellular compartments (Fig 2). However, SAHH1 was not uniformly localized within the cells, but rather highly organized to various cellular structures. SAHH1 was found dynamically associated with cytoplasmic strands, along the plasma membrane, in punctate structures, and around chloroplasts (Fig 2, S1 Video). In line with a previous report by Lee et al. (2012), strong fluorescence arising from EGFP-SAHH1 was also detected in the nucleus (Fig 2E). The nucleolus, however, was completely devoid of SAHH1 (Fig 2E). Immunoblot analysis of protein extracts separated on Clear Native (CN) gels revealed the presence of EGFP-SAHH1 in oligomeric protein complexes, and particularly the abundant SAHH complex 4, as was observed in wild type (Fig 2G) [17]. Moreover, immunoblot analysis of SDS-gels confirmed the presence of EGFP-SAHH1 in the leaf extracts, whereas free EGFP could not be observed (S1 Fig).

**Fig 2.**
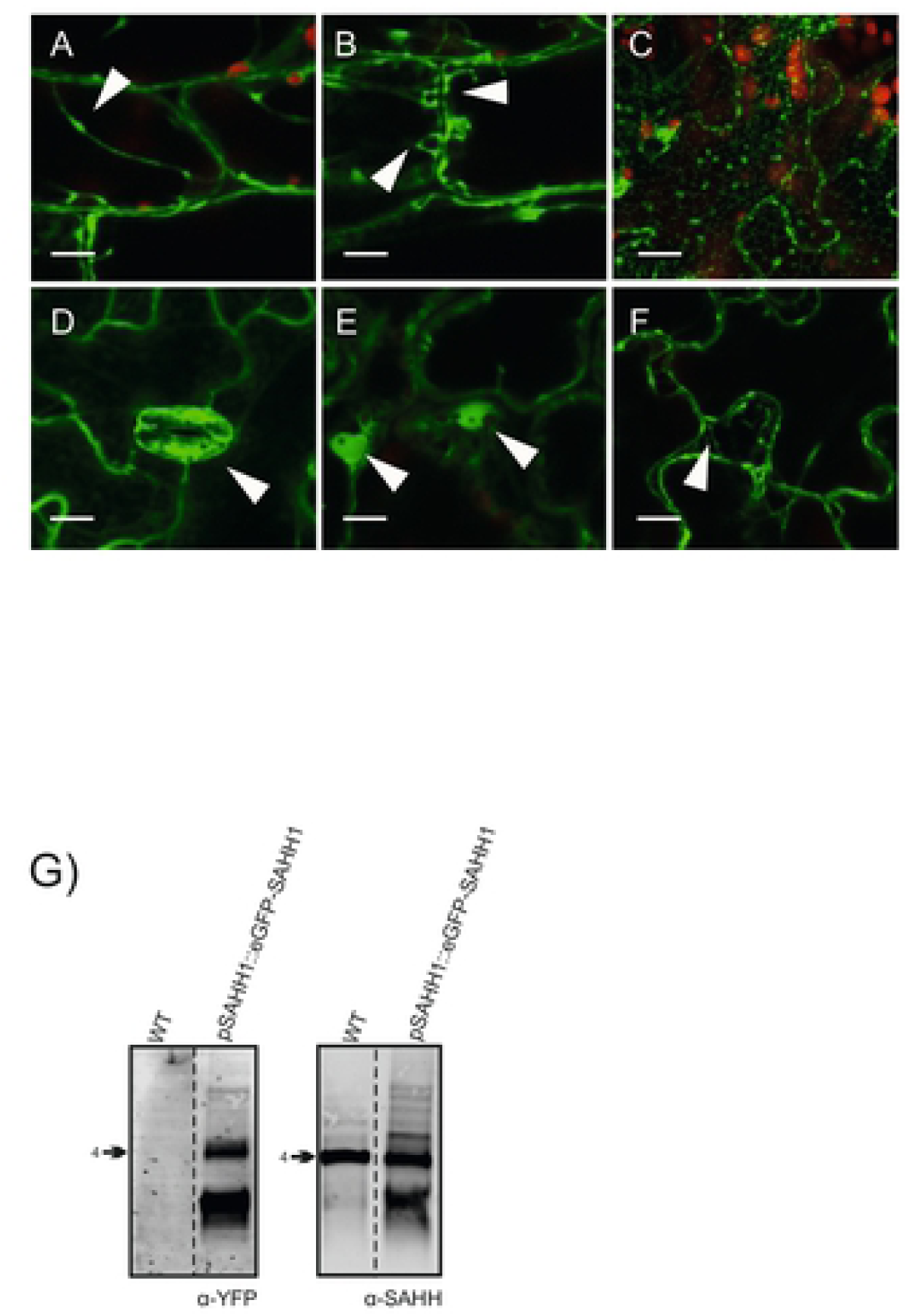
Sub-cellular localization of SAHH1 in Arabidopsis leaves. A-F) Confocal microscopy images obtained from transgenic plants stably expressing SAHH1p::EGFP-SAHH1. EGFP-SAHH1 was localized to cytoplasmic strands with associated bodies (A), vesicular structures (B), reticulate constructions (C), stomatal guard cells (D), nuclei (E) and the plasma membrane (F). The red color corresponds to chlorophyll autofluorescence from chloroplasts. The scale bars correspond to10 µm. G) Immunoblot analysis depicting the presence of EGFP-SAHH1 in oligomeric protein complexes. Leaf extracts from Arabidopsis Col-0 and a transgenic line expressing SAHH1p::EGFP-SAHH1 were isolated, separated on Clear Native gel electrophoresis and immunodetected with anti-SAHH antibody.

### Assessment of Arabidopsis SAHH by 2D gel electrophoresis

Previously, we reported the presence of SAHH in multiple oligomeric compositions and the co-migration of the abundant SAHH complex 4 with another abundant protein spot containing e.g. CARBONIC ANHYDRASE 1 (CA1; previously identified as the chloroplastic SALICYLIC ACID-BINDING PROTEIN 1 SABP3) [26], on a 3D Clear Native gel system [17]. The following proteomic approach was therefore designed to decipher whether CA1 is a component in the SAHH complex 4 (Fig 3).

**Fig 3.**
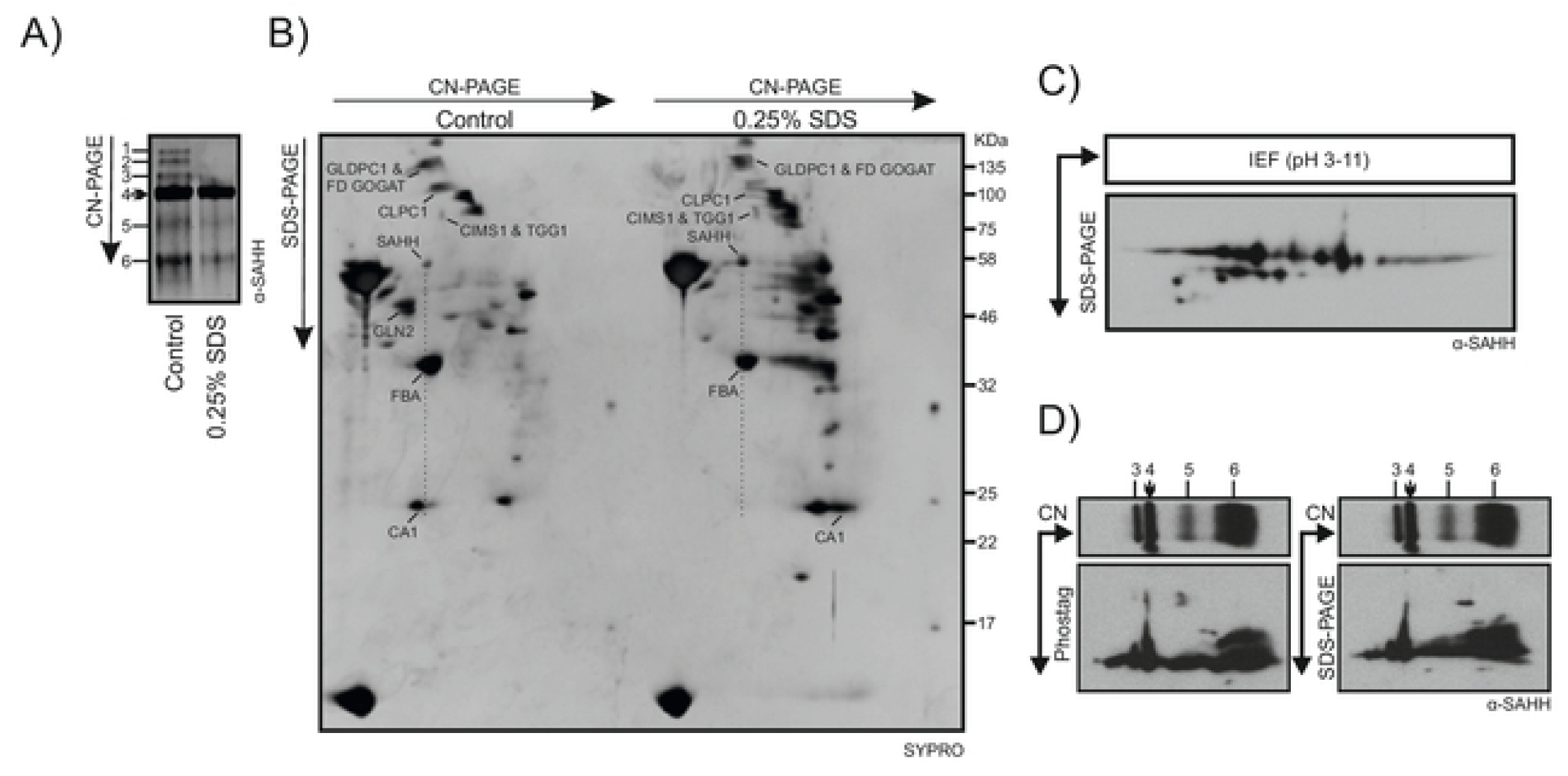
Biochemical analysis of Arabidopsis SAHH complex 4. A) Arabidopsis leaf extracts were incubated in the presence and absence of 0.25% SDS and thereafter separated on Clear Native (CN) gels followed by 12% acrylamide SDS-PAGE in the second dimension. 1D CN-PAGE immunoblot with anti-SAHH is shown on the top of a 2D SDS-PAGE, which was stained with a total protein stain (SYPRO). The SAHH-containing major spots originating from the SAHH complex 4, CARBONIC ANHYDRASE 1 (CA1), FRUCTOSE BISPHOSPHATE ALDOLASE (FBA), GLYCINE DECARBOXYLASE P-PROTEIN 1 (GLDPC1), FERREDOXIN-DEPENDENT GLUTAMATE SYNTHASE 1 (FD GOGAT), CLPC HOMOLOGUE 1 (CLPC1), COBALAMIN-INDEPENDENT METHIONINE SYNTHASE (CIMS1), THIOGLUCOSIDE GLUCOHYDROLASE 1 (TGG1), and GLUTAMINE SYNTHASE (GLN) are marked. B) Arabidopsis leaf extracts were fractionated by isoelectric focusing in a pH range from 3 to 11 and subsequently separated by 12% acrylamide SDS-PAGE in the second dimension. C) Arabidopsis leaf extracts were fractionated on CN-PAGE and the protein complexes were thereafter separated on 7.5% acrylamide Phostag-PAGE (left panel) or on 7.5% acrylamide Phostag-PAGE lacking the Phostag reagent (right panel).

The stability of the SAHH-containing complexes was first assessed using SDS as a detergent and DTT as a reducing agent. Upon treatment of soluble leaf extracts with 1% SDS, the SAHH complexes 1, 2 and 3 were disrupted and the SAHH complexes 4 and 5 became less abundant when separated by CN-PAGE (S2 Fig). In contrast, treatment of soluble leaf extracts with 10 mM DTT did not affect the stability of SAHH complex 4 (S2 Fig).

Immunoblot analysis with anti-SAHH antibody suggested that pre-treatment of soluble leaf extracts with 0.25% SDS did not alter the migration of SAHH complex 4 on CN-PAGE although it tore GLN2 complex (Fig 3A) showing the stability of protein complexes was differentially affected by the 0.25% SDS treatment. However, when the protein complexes were separated on 2D CN-PAGE, the abundant CA1-containing protein spot disappeared in SDS-treated samples, indicating that the spot did not contain stoichiometric components of SAHH complex 4 (Fig 3A). The CA1 shift on the 2D SDS-PAGE is due to monomerization of the complex by the treatment. Another abundant protein spot that co-migrated with SAHH complex 4 was identified as chloroplastic Fructose Bisphosphate Aldolase (FBA) [17], which does not co-localize with SAHH (Fig 2A) and is therefore highly unlikely to interact with SAHH. Based on these findings and the apparent 200 kDa MW of the complex, it can be deduced that the SAHH complex 4 is composed by a tetramer of the enzyme. Shattered of complexed 1 and 2 with 0.25% treatment generate the increase in abundance of SAHH complex 4 containing spot in the 0.25% SDS treatment (Fig 3 A and B).

Next we assessed the extent to which SAHH is present in different forms in Arabidopsis leaf extracts. Isoelectric focusing and 2D SDS-PAGE, followed by immunoblot analysis detected SAHH in multiple spots with different pIs and three different molecular masses (Fig 3B). Such variety of combinations may arise from various combinations of PTMs, which could allow enormous versatility in the regulation of SAHH function.

The final approach was designed to explore whether SAHH is differentially phosphorylated within the different oligomeric compositions. To this end, soluble leaf extracts were run on CN-PAGE, followed by separation of differentially phosphorylated proteins on Phostag gels in the second dimension (Fig 3C). In parallel, a control sample of equal protein content was separated on an SDS-PAGE devoid of urea. Immunoblotting of the Phostag gels with anti-SAHH antibody revealed slow-migrating protein spots, indicative of SAHH phosphorylation in complexes 3, 4 and 5 (Fig 3C).

### Evolutionary conservation of SAHH in land plants

To gain insights into evolutionary conservation of SAHH, we first compared the amino acid sequences of SAHH in divergent plant species, including *A. thaliana, L. luteus, B. oleracea, S. oleracea* and *P. patens* (Fig 4). Arabidopsis and *B. oleracea* are closely related species and showed 99% SAHH amino acid sequence similarity (Fig 4 A and B, Table 1). Even between the more distantly related species, pair-wise amino acid comparison between *L. luteus* and Physcomitrella SAHH indicated 90% similarity (Fig 4 A and B, Table 1). Arabidopsis SAHH1 has been reported to undergo phosphorylation [17,27–30], S-nitrosylation [31,32], acetylation [28] and ubiquitination [33] at multiple sites. The majority of the experimentally described PTMs sites were conserved in the plant species studied (Fig 4A), suggesting that post-translational regulation of SAHH could be a conserved feature among land plants (Fig 4A).

**Table 1.** Pair wise comparison of SAHH1 amino acid sequences from *A. thaliana, L. luteus, B. oleracea, S. oleracea* and *P. patens*. Identities and similarities are shown.

|  | IDENTITIES | SIMILARITIES |
| --- | --- | --- |
| *Arabidopsis* & *L. luteus* | 0,9016393 | 0,9487705 |
| *Arabidopsis* & *B. oleracea* | **0,9670103** | **0,9917526** |
| *Arabidopsis* & *S. oleracea* | 0,8870637 | 0,9178645 |
| *Arabidopsis* & *P. patens* | 0,8721649 | 0,9319588 |
| *L. luteus* & *B. oleracea* | 0,8954918 | 0,9467213 |
| *L. luteus* & *S. oleracea* | 0,8750000 | 0,9159836 |
| *L. luteus* & *P. patens* | **0,8463115** | **0,9036885** |
| *B. oleracea* & *S. oleracea* | 0,8850103 | 0,9445585 |
| *B. oleracea* & *P. patens* | 0,8680412 | 0,9360825 |
| *S. oleracea* & *P. patens* | 0,8542094 | 0,9178645 |

**Fig 4.**
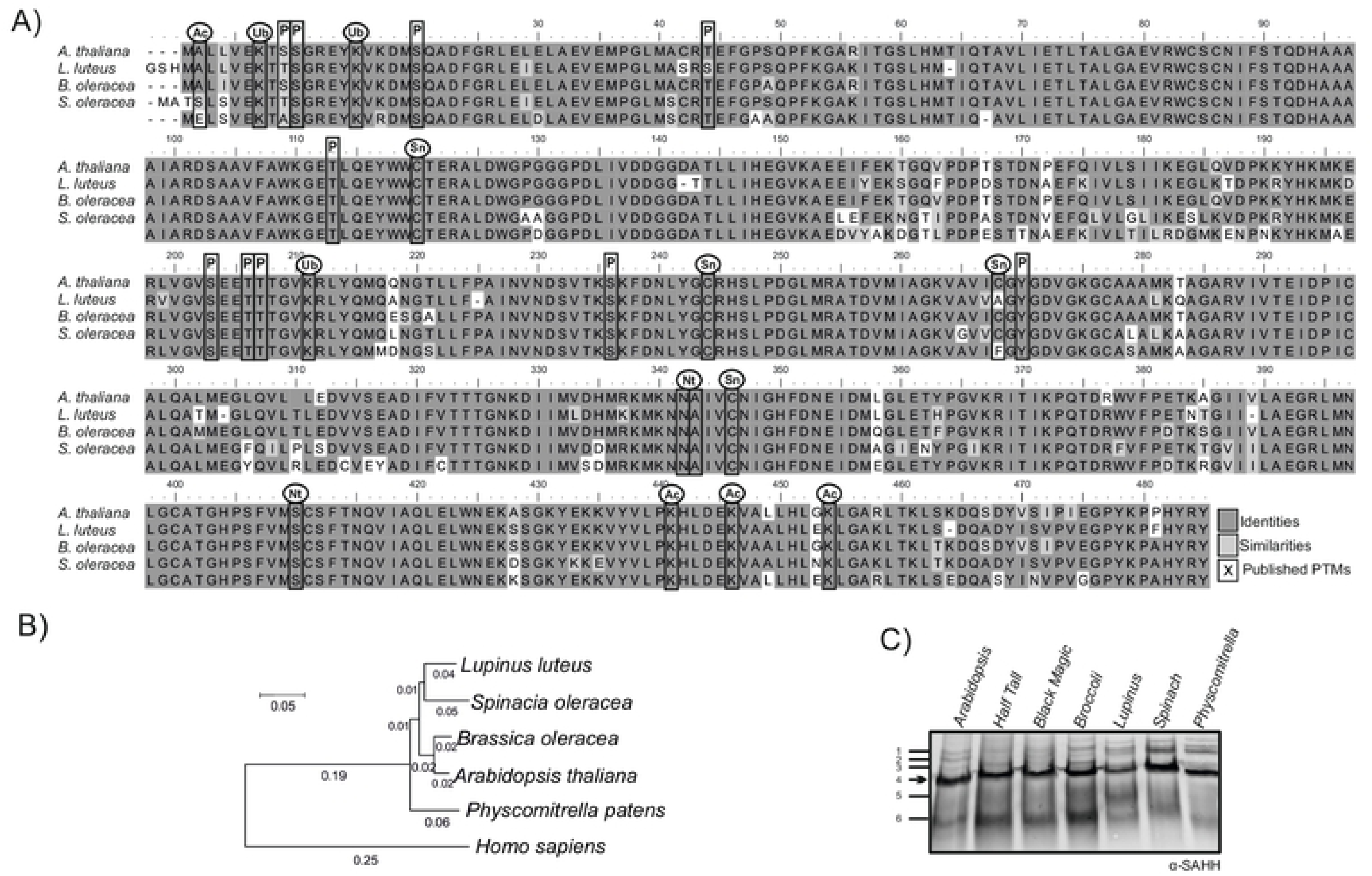
Evolutionary conservation of SAHH on protein level. A) Amino acid sequence alignment of SAHH1 between *A. thaliana, L. luteus, B. oleracea, S. oleracea* and *P. patens*. Ac, Lysine Acetylation; P, phosphorylation; Nt, N-terminus Proteolysis; Sn, S-nitrosylation; Ub, ubiquitination. B) Phylogenetic tree based on SAHH1 amino acid sequence constructed using the neighbor-joining method. Numbers represent substitution per amino acid based on data from 500 trees. C) SAHH containing protein complexes in evolutionarily divergent plant species. Six SAHH containing protein complexes as detected by anti-SAHH antibody in total soluble protein extracts of *A. thaliana, B. oleracea* convar *italic* (broccoli), *B. oleracea* convar *acephala* var. Half Tall (kale) and *B. oleracea* convar *acephala* Black Magic (kale), *L. luteus, S. oleracea* and *P. patens* separated on CN-PAGE. The abundant SAHH-containing protein complex is indicated by arrow.

Next we assessed the conservation of SAHH complex formation across evolutionarily distant plants by CN-PAGE and immunoblotting with anti-SAHH antibody. This approach revealed presence of the predominant SAHH complex 4 in all the plant species studied (Fig 4C). The presence of SAHH complex 4 was also detected in protein extracts isolated from *L. luteus*, for which the resolved crystal structure suggested that SAHH would be active as a dimer [14,15]. Moreover, all the species studied also displayed other SAHH complexes that migrated similarly to those observed in Arabidopsis when separated by CN-PAGE. Taken together, these results suggested that SAHH is a conserved enzyme, which forms similar oligomeric protein complexes in phylogenetically different land plants.

### Light-induced adjustments of SAHH in Arabidopsis and Physcomitrella

To assess stress-induced adjustments in SAHH, we next exposed the two well-established, phylogenetically different model plants, Arabidopsis and Physcomitrella, to high irradiance levels for two days. Separation of SAHH-containing complexes on CN-PAGE revealed a subtle high-light-induced decrease in SAHH complex 2 abundance, and a corresponding increase in the abundance of SAHH complexes 5 and 6 in Arabidopsis (Fig 5A). Physcomitrella, in contrast, showed a clear increase in the abundance of a SAHH complex, which corresponds to SAHH complex 2 in Arabidopsis (Fig 5A). The total abundance of SAHH did not differ between the treatments (Fig 5C). The high light treatment slightly decreased SAHH complex 4 abundance in Arabidopsis, whereas Physcomitrella accumulated slightly more SAHH complex 4 (Fig 5A). Hence, both model species responded to high light treatment at the level of SAHH complex formation, but exhibited opposite outcomes.

**Fig 5.**
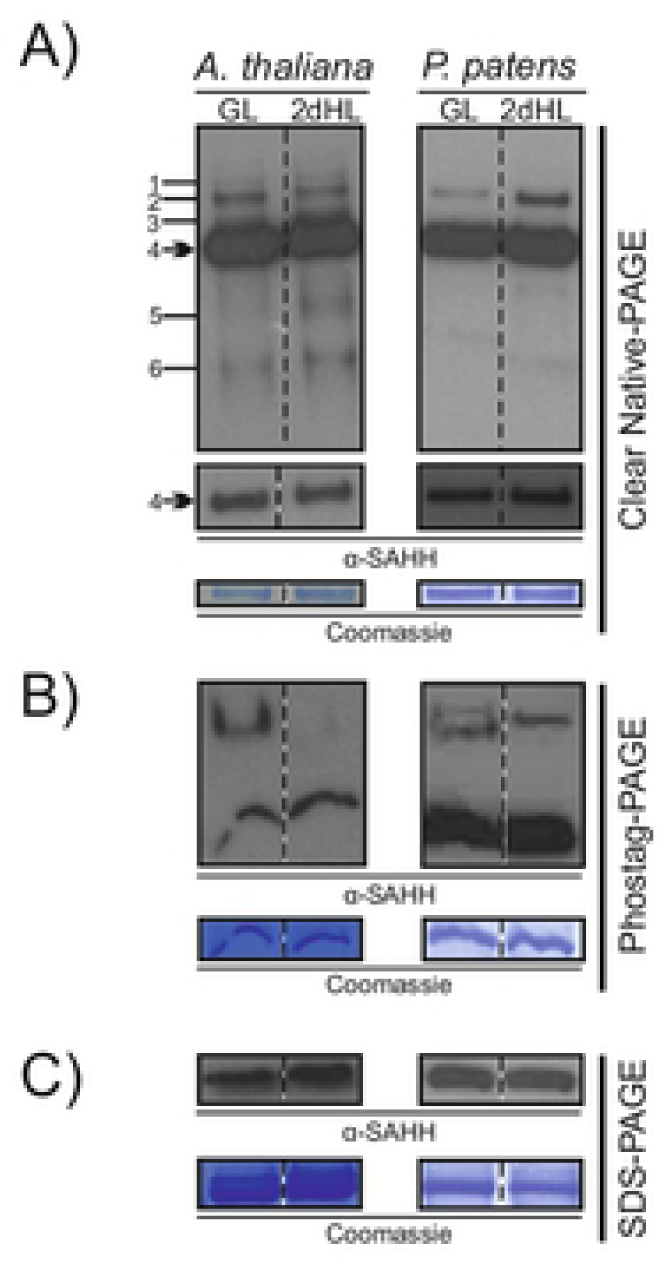
Light-stress-induced adjustments in SAHH. *Arabidopsis thaliana* was grown under 130 µmol photons m^-2^ s^-1^ for 16 days and thereafter shifted 800 µmol photons m^-2^ s^-1^ for 2 days. *Pyscomitrella patens* as a shade-adapted moss species was grown under 45 µmol photons m^-2^ sec^-1^ for 13 days and thereafter illuminated under 500 µmol photons m^-2^ s^-1^ for two days. SAHH was separated by gel-based systems and immunodetected by using an anti-SAHH antibody. Coomassie-stained membranes are shown as load controls for each experiment. A) Oligomeric SAHH complexes as detected by clear native (CN)-PAGE in Arabidopsis and Pyscomitrella in growth light (GL) and after 2-day illumination under high light (2dHL). Six oligomeric SAHH complexes are indicated by numbers. The lower panel depicts an immunoblot with a shorter exposure time for visualization of the abundant SAHH complex 4. B) SAHH protein phosphorylation as detected by Phostag-PAGE in Arabidopsis and Pyscomitrella in growth light (GL) and after 2-day illumination under high light (2dHL). C) SAHH protein abundance as detected by SDS-PAGE in Arabidopsis and Pyscomitrella in growth light (GL) and after 2-day illumination under high light (2dHL).

Analysis of SAHH phosphorylation by Phos-tag gel electrophoresis suggested also light-dependent phosphoregulation in both species (Fig 5B). In Arabidopsis, a slow-migrating form of SAHH was detected in leaf extracts isolated from growth light conditions, while high-light-exposed leaves did not contain such phosphorylated form of SAHH (Fig 5B). In Physcomitrella, the intensity of one phosphorylated SAHH species increased, while another SAHH species disappeared upon high light illumination (Fig 5B). Hence, both Arabidopsis and Physcomitrella responded to light-induced stress by regulatory adjustments in SAHH, but the responses differed between the angiosperm and bryophyte models.

## DISCUSSION

### SAHH is required to maintain trans-methylation reactions in different cellular compartments

Trans-methylation reactions are intrinsic to cellular metabolism and a prerequisite for normal plant growth and development. Reflecting the enormous diversity of species-specific trans-methylation reactions that take place in metabolic and regulatory networks, adjustments in SAHH function can be expected to differ under different physiological states in different species. SAM-dependent trans-methylation reactions occur in a multitude of subcellular compartments where SAH must be efficiently metabolized to maintain MT activities [4]. Functional characterization of Arabidopsis mutants has demonstrated that loss of SAHH function can result in global inhibition of MT activities because of accumulation of SAH [34]. Changes in the sub-cellular localization of SAHH could also significantly affect the efficiency of specific trans-methylation reactions [18,35,36]. Associated with this, SAHH can translocate from the cytoplasm into the nucleus [18] and is also dynamically distributed in other sub-cellular compartments, including vesicular structures, reticulate constructions and the plasma membrane, but not chloroplasts or mitochondria (Fig 2).

One of the key functions of SAHH is to maintain appropriate patterns of DNA and histone methylation in the nucleus [6,35]. The nuclear localization of SAHH was recently attributed to a 41-amino-acid segment (Gly150-Lys190), which is a prerequisite for nuclear targeting of Arabidopsis SAHH1 [18]. Intriguingly, the surface-exposed segment does not act as an autonomous nuclear localization signal *per se*, but may rather serve as an interaction domain for associations with other proteins that can direct SAHH into the nucleus. In line with this idea, it was proposed that physical interactions between SAHH and MTs could provide a means for targeting SAHH to appropriate subcellular compartments in order to ensure uninterrupted trans-methylation [18].

While SAHH1 has not been localized into the chloroplast (Fig 2) [18], we detected SAHH1 as a ring in the immediate vicinity around the photosynthetic organelles (Fig 2), presumably to facilitate efficient removal of SAH upon export by SAM transporters that exchange SAH for SAM synthesized in the cytoplasm [37,38]. We also found that SAHH1 can dynamically move along cytosolic strands (S1 Video), but the possible mechanisms underlying such sub-cellular movements remain to be established. Besides protein interactions with MTs, physical contact with components of the cytoskeleton or enzymatic protein complexes may direct SAHH to appropriate sub-cellular localizations.

SAHH forms oligomeric complexes and undergoes dynamic interactions with various endogenous and exogenous proteins [2,39,40]. SAHH has been shown to physically interact with various methyltransferases, including indole glucosinolate methyltransferases [17], mRNA cap methyl-transferase [18], and caffeoyl CoA methyltransferase (CCoAOMT) [36], presumably to ensure efficient trans-methylation reactions at accurate sub-cellular sites. These interactions are likely largely determined by the availability of MTs, which appear to be transcriptionally highly responsive to endogenous and exogenous cues. Yang et al. (2019) proposed that *in vivo* interactions between CCoAOMT7, SAHH and SAMS form a complex for SAM synthesis to enhance the formation of ferulate in the cell wall. Interactions between SAHH and other proteins may also be affected by different combinations of post-translational modifications.

Based on crystallographic studies, plant SAHH is proposed to be active as a dimer [14,15]. However, our data provide evidence suggesting that in Arabidopsis SAHH is predominantly present as a tetramer. This was evidenced by treatment of leaf extracts with a low concentration of SDS, which did not affect the presence of SAHH complex 4 on CN gels, but abolished co-migration of potential complex-forming proteins when assessed by 2D SDS-PAGE (Fig 3A). The physiological significance of this abundant SAHH complex present in various land plants, including the moss Physcomitrella (Fig 4C), remains to be established. Likewise, the subunit composition of the other SAHH complexes reported here remains to be uncovered. Moreover, besides formation of biochemically rather stable oligomeric complexes, SAHH is likely to undergo transient interactions that cannot be trapped by physical separation of protein complexes.

### SAHH is an evolutionary conserved enzyme governed by multilevel post-translational control

The metabolic centrality of SAHH is reflected by its evolutionary conservation and the high number of PTMs, including phosphorylation, S-nitrosylation, acetylation and ubiquitination, which have been experimentally verified to occur on Arabidopsis SAHH [17,28–33,41]. Conservation of the PTM sites between Physcomitrella, a brypohyte, and angiosperms (Fig 4A) points to multilevel post-translational regulation that can facilitate delicate metabolic responses to environmental cues. The up-stream regulatory enzymes, such as the protein kinases and protein phosphatases, N-acetyl transferases and ubiquitin ligases, however, remain almost completely unidentified. Hints to phosphoregulation of SAHH were provided by Trotta et al. [23] and Rahikainen et al. [17], who provided evidence that a protein phosphatase 2A regulatory subunit PP2A-B′γ controls SAHH complex formation and the associated trans-methylation capacity of leaf cells. Two of the phosphorylated residues on Arabidopsis SAHH1, S203 and S236, reside on conserved amino acids in the active center of the enzyme [15], and phosphorylation at these sites could therefore affect the activation state of the enzyme. The phosphorylated residues S20 and T44 of SAHH1 in turn reside on the surface of the enzyme [15], and changes in phosphorylation of these sites could impact its protein interactions and/or sub-cellular localization.

Whether and how the different functional aspects of SAHH respond to environmental signals in different plant species is a key question to be resolved to understand metabolic regulation in plants. Here, we explored how the biochemical characteristics of SAHH become adjusted in response to light stress, which poses a risk of metabolic imbalance and photo-oxidation in photosynthetic tissues. To avoid light-induced damage, angiosperms undergo coordinated acts to balance the function of photosynthetic electron transfer and the down-stream carbon metabolism [42–44]. More recently, studies on Physcomitrella have revealed the existence of multilayered mechanisms that jointly protect the shade-acclimated bryophyte against photoinhibition [45].

Excess light also triggers chloroplast-to-nucleus signaling, whereby stress-exposed chloroplasts induce coordinated adjustments in the expression of nuclear genes [46,47]. Light-induced metabolic adjustments, beyond photosynthetic carbon metabolism and other chloroplastic pathways, in turn remain poorly understood. Recently, Zhao et al. (2019) demonstrated that a conserved signaling mechanism, where a chloroplast retrograde signal interacts with hormonal signaling to drive stomatal closure, is operational in angiosperms, mosses and ferns [48]. Our studies provide evidence that high-light-induced signals, likely originating from chloroplasts, are reflected by regulatory adjustments in SAHH at the level of complex formation and phosphorylation in both Physcomitrella and Arabidopsis (Fig 5A).

Taken together, cellular trans-methylation reactions are largely determined by the abundance of substrate-specific MTs that require the activated methyl cycle to retain their activity. Among AMC enzymes, the functionality of SAHH is controlled at the level of sub-cellular localization, complex formation and post-translational modifications, which can modulate the activity and interactions between SAHH and other proteins. Jointly, these regulatory actions determine which MTs get to interact with SAHH, thereby maintaining their activity. Hence, SAHH can be considered a key determinant of trans-methylation reactions in living cells.

## Acknowledgements

We thank Marianna Alaviuhkola for excellent assistance in microscopy. The confocal imaging was performed with microscopes of the Cell Imaging and Cytometry Core at the Turku Bioscience Centre, University of Turku and Åbo Akademi University. We are grateful to Dr. Caterina Gerotto for providing *Pyscomitrella patens* material. This work was financially supported by Academy of Finland project 307719 to SK, 325122 to JP, and the Academy of Finland Center of Excellence in Primary Producers 2014-2019 (307335). SA and MR were funded by the University of Turku Doctoral Programme in Molecular Life Sciences, the Turku University Foundation and the Finnish Cultural Foundation Varsinais-Suomi Regional Fund. MB was funded by the University of Helsinki.

## SUPPORTING INFORMATION

**S1 Table. List of reviewed *Arabidopsis thaliana* O-methyltransferases retrieved from UniProt and AMC enzymes used as input for GENESTIGATOR analysis.** Clusters according to the performed hierarchical cluster analysis, protein accession, entry and protein name are indicated. Activated methyl cycle enzymes (AMC) are marked in blue.

**S1 Fig. Immunoblots depicting EGFP-SAHH1 in *Arabidopsis thaliana* wild type (WT) and a transgenic line stably expressing SAHH1p::EGFP-SAHH1**. Proteins were separated on SDS-PAGE, and EGFP-SAHH1 was immunodetected with an anti-YFP antibody and SAHH was detected with an anti-SAHH antibody.

**S2 Fig. Immunoblot depicting SAHH protein complexes after treatment of *Arabidopsis thaliana* foliar leaf extracts with SDS and/or DTT.** For combined treatments with SDS and DTT, the leaf extract was incubated in the presence of one chemical for 30 minutes, followed by addition of the other for 30 minutes.

**S1 Video. Dynamic movements of SAHH1p::EGFP-SAHH1 in Arabidopsis cells.**

## REFERENCES

1. Moffatt B, Weretilnyk E. Sustaining S-adenosyl-L-methionine-dependent methyltransferase activity in plant cells. Physiol Plant [Internet]. 2001;113(4):435–42. Available from: http://www.blackwell-synergy.com/links/doi/10.1034/j.1399-3054.2001.1130401.x

2. Rahikainen M, Alegre S, Trotta A, Pascual J, Kangasjärvi S. Trans-methylation reactions in plants: focus on the activated methyl cycle. Physiol Plant. 2018;162(2):162–76.

3. Richards HH, Chiang PK, Cantoni GL. Adenosylhomocysteine hydrolase: Crystalizationof the purified enzyme and its properties. J Biol Chem. 1978;253:4476–4480.

4. Poulton JE. Transmethylation and Demethylation Reactions in the Metabolism of Secondary Plant Products. In: Secondary Plant Products [Internet]. Academic Press; 1981 [cited 2019 Dec 2]. p. 667–723. Available from: https://www.sciencedirect.com/science/article/pii/B9780126754070500282?via%3Dihub

5. Cantoni GL, Scarano E. The formation of s-adenosylhomocysteine in enzymatic transmethylation reactions. J Am Chem Soc. 1954;76(18):4744.

6. Rocha PSCF, Sheikh M, Melchiorre R, Fagard M, Boutet S, Loach R, et al. The arabidopsis HOMOLOGY-DEPENDENT GENE SILENCING1 gene codes for an S-adenosyl-L-homocysteine hydrolase required for DNA methylation-dependent gene silencing. Plant Cell. 2005;17(2):404–17.

7. Pereira LAR, Todorova M, Cai X, Makaroff CA, Emery RJN, Moffatt BA. Methyl recycling activities are co-ordinately regulated during plant development. J Exp Bot. 2007;58(5):1083–98.

8. Li CH, Yu N, Jiang SM, Shangguan XX, Wang LJ, Chen XY. Down-regulation of S-adenosyl-L-homocysteine hydrolase reveals a role of cytokinin in promoting transmethylation reactions. Planta. 2008;228(1):125–36.

9. Barić I, Fumić K, Glenn B, Ćuk M, Schulze A, Finkelstein JD, et al. S-adenosylhomocysteine hydrolase deficiency in a human: A genetic disorder of methionine metabolism. Proc Natl Acad Sci U S A. 2004;101(12):4234–9.

10. Barić I, Ćuk M, Fumić K, Vugrek O, Allen RH, Glenn B, et al. S-Adenosylhomocysteine hydrolase deficiency: A second patient, the younger brother of the index patient, and outcomes during therapy. J Inherit Metab Dis. 2005;28(6):885–902.

11. Wu X, Li F, Kolenovsky A, Caplan A, Cui Y, Cutler A, et al. A mutant deficient in S-adenosylhomocysteine hydrolase in Arabidopsis shows defects in roothair development. Botany. 2009;87(6):571–84.

12. Kusakabe Y, Ishihara M, Umeda T, Kuroda D, Nakanishi M, Kitade Y, et al. Structural insights into the reaction mechanism of S-adenosyl-L-homocysteine hydrolase. Sci Rep [Internet]. 2015;5:1–16. Available from: http://dx.doi.org/10.1038/srep16641

13. Turner MA, Yuan C, Borchardt RT, Hershfield MS, Smith GD, Howell PL. Structure determination of selenomethionyl S-adenosylhomocysteine hydrolase using data at a single wavelength. 1998;5(5):369–76.

14. Brzezinski K, Bujacz G, Jaskolski M. Purification, crystallization and preliminary crystallographic studies of plant S -adenosyl-L -homocysteine hydrolase (*Lupinus luteus*). Acta Crystallogr. 2008;64:671–3.

15. Brzezinski K, Dauter Z, Jaskolski M. High-resolution structures of complexes of plant S -adenosyl-L -homocysteine hydrolase (*Lupinus luteus*). Acta Crystallogr. 2012;D68:218–31.

16. Matuszewska B, Borchardt RT, Muto N, Tsujino M, Sudate Y, Hayashi M, et al. The Mechanism of Inhibition of Alcaligenes faecalis Hydrolase by Neplanocin A. 1987;256(1):50–5.

17. Rahikainen M, Trotta A, Alegre S, Pascual J, Vuorinen K, Overmyer K, et al. PP2A-B’γ modulates foliar trans -methylation capacity and the formation of 4-methoxy-indol-3-yl-methyl glucosinolate in Arabidopsis leaves. Plant J. 2017;89:112–27.

18. Lee S, Doxey AC, Mcconkey BJ, Moffatt BA. Nuclear Targeting of Methyl-Recycling Enzymes in *Arabidopsis thaliana* Is Mediated by Specific Protein Interactions. Mol Plant [Internet]. 2012;5(1):231–48. Available from: http://dx.doi.org/10.1093/mp/ssr083

19. Gerotto C, Trotta A, Bajwa AA, Mancini I, Morosinotto T, Aro EM. Thylakoid protein phosphorylation dynamics in a moss mutant lacking SERINE/THREONINE PROTEIN KINASE STN8. Plant Physiol. 2019;180(3):1582–97.

20. Hruz T, Laule O, Szabo G, Wessendorp F, Bleuler S, Oertle L, et al. Genevestigator V3: A Reference Expression Database for the Meta-Analysis of Transcriptomes. Adv Bioinformatics [Internet]. 2008;2008:1–5. Available from: http://www.hindawi.com/journals/abi/2008/420747/

21. Kolde R. pheatmap: Pretty Heatmaps. R package version 1.0. 12. 2019.

22. Lehtimäki N, Koskela MM, Dahlström KM, Pakula E, Lintala M, Scholz M, et al. Posttranslational modifications of FERREDOXIN-NADP+ OXIDOREDUCTASE in arabidopsis chloroplasts. Plant Physiol. 2014;166(4):1764–76.

23. Trotta A, Wrzaczek M, Scharte J, Tikkanen M, Konert G, Rahikainen M, et al. Regulatory subunit B’gamma of protein phosphatase 2A prevents unnecessary defense reactions under low light in Arabidopsis. Plant Physiol. 2011;156(3):1464–80.

24. Thompson JD, Higgins DG, Gibson TJ. CLUSTAL W: Improving the sensitivity of progressive multiple sequence alignment through sequence weighting, position-specific gap penalties and weight matrix choice. Nucleic Acids Res. 1994;22(22):4673–80.

25. Saitou N, Nei M. The neighbor-joining method: a new method for reconstructing phylogenetic trees. Mol Biol Evol. 1987;4(4):406–25.

26. Slaymaker DH, Navarre DA, Clark D, Del Pozo O, Martin GB, Klessig DF. The tobacco salicylic acid-binding protein 3 (SABP3) is the chloroplast carbonic anhydrase, which exhibits antioxidant activity and plays a role in the hypersensitive defense response. Proc Natl Acad Sci U S A. 2002;99(18):11640–5.

27. Li L, Li M, Yu L, Zhou Z, Liang X, Liu Z, et al. The FLS2-Associated Kinase BIK1 Directly Phosphorylates the NADPH Oxidase RbohD to Control Plant Immunity. Cell Host Microbe. 2014;15(3):329–38.

28. Xu SL, Chalkley RJ, Maynard JC, Wang W, Ni W, Jiang X, et al. Proteomic analysis reveals O-GlcNAc modification on proteins with key regulatory functions in Arabidopsis. PNAS. 2017;114(8):E1536–43.

29. Roitinger E, Hofer M, Ko T, Pichler P, Novatchkova M, Yang J, et al. Quantitative Phosphoproteomics of the Ataxia Telangiectasia-Mutated (ATM) and Ataxia Telangiectasia-Mutated and Rad3-related (ATR) Dependent DNA Damage Response in *Arabidopsis thaliana*. Mol Cell proteomics. 2015;556–71.

30. Wang X, Bian Y, Cheng K, Gu L, Ye M, Zou H, et al. A large-scale protein phosphorylation analysis reveals novel phosphorylation motifs and phosphoregulatory networks in Arabidopsis. J Proteomics [Internet]. 2013;78:486–98. Available from: http://dx.doi.org/10.1016/j.jprot.2012.10.018

31. Fares A, Rossignol M, Peltier J. Biochemical and Biophysical Research Communications Proteomics investigation of endogenous S -nitrosylation in Arabidopsis. Biochem Biophys Res Commun [Internet]. 2011;416(3–4):331–6. Available from: http://dx.doi.org/10.1016/j.bbrc.2011.11.036

32. Puyaubert J, Fares A, Rézé N, Peltier J, Baudouin E. Plant Science Identification of endogenously S-nitrosylated proteins in Arabidopsis plantlets: Effect of cold stress on cysteine nitrosylation level. Plant Sci [Internet]. 2014;215–216:150–6. Available from: http://dx.doi.org/10.1016/j.plantsci.2013.10.014

33. Walton A, Stes E, Cybulski N, Bel M, Iñigo S, Durand AN, et al. It’s time for some “site”-seeing: Novel tools to monitor the ubiquitin landscape in *Arabidopsis thaliana*. Plant Cell. 2016;28(1):6–16.

34. Jordan ND, West JP, Bottley A, Sheikh M, Furner I. Transcript profiling of the hypomethylated *hog1* mutant of Arabidopsis. Plant Mol Biol. 2007;65(5):571–86.

35. Baubec T, Dinh HQ, Pecinka A, Rakic B, Rozhon W, Wohlrab B, et al. Cooperation of multiple chromatin modifications can generate unanticipated stability of epigenetic states in Arabidopsis. Plant Cell. 2010;22(1):34–47.

36. Yang SX, Wu TT, Ding CH, Zhou PC, Chen ZZ, Gou JY. SAHH and SAMS form a methyl donor complex with CCoAOMT7 for methylation of phenolic compounds. Biochem Biophys Res Commun [Internet]. 2019;520(1):122–7. Available from: https://doi.org/10.1016/j.bbrc.2019.09.101

37. Ravanel S, Block MA, Rippert P, Jabrin S, Curien G, Rébeillé F, et al. Methionine metabolism in plants: Chloroplasts are autonomous for de novo methionine synthesis and can import S-adenosylmethionine from the cytosol. J Biol Chem. 2004;279(21):22548–57.

38. Bouvier F, Linka N, Isner JC, Mutterer J, Weber APM, Camara B. Arabidopsis SAMT1 defines a plastid transporter regulating plastid biogenesis and plant development. Plant Cell. 2006;18(11):3088–105.

39. Mäkinen K, De S. The significance of methionine cycle enzymes in plant virus infections. Curr Opin Plant Biol. 2019;50:67–75.

40. Ivanov KI, Eskelin K, Bašic M, De S, Lõhmus A, Varjosalo M, et al. Molecular insights into the function of the viral RNA silencing suppressor HCPro. Plant J. 2016;85(1):30–45.

41. Li S, Mhamdi A, Trotta A, Kangasjärvi S, Noctor G. The protein phosphatase subunit PP2A-B’γ is required to suppress day length-dependent pathogenesis responses triggered by intracellular oxidative stress. New Phytol. 2014;202(1):145–60.

42. Aro EM, Virgin I, Andersson B. Photoinhibition of Photosystem II. Inactivation, protein damage and turnover. BBA - Bioenerg. 1993;1143(2):113–34.

43. Foyer CH. Reactive oxygen species, oxidative signaling and the regulation of photosynthesis. Environ Exp Bot [Internet]. 2018;154(May):134–42. Available from: https://doi.org/10.1016/j.envexpbot.2018.05.003

44. Pascual J, Rahikainen M, Kangasjärvi S. Plant Light Stress. eLS. 2017;(April):1–6.

45. Alboresia A, Gerottob C, Giacomettib GM, Bassia R, Morosinotto T. Physcomitrella patens mutants affected on heat dissipation clarify the evolution of photoprotection mechanisms upon land colonization. Proc Natl Acad Sci U S A. 2010;107(24):11128–33.

46. Kleine T, Leister D. Retrograde signaling: Organelles go networking. Biochim Biophys Acta - Bioenerg [Internet]. 2016;1857(8):1313–25. Available from: http://dx.doi.org/10.1016/j.bbabio.2016.03.017

47. Chan KX, Phua SY, Crisp P, McQuinn R, Pogson BJ. Learning the Languages of the Chloroplast: Retrograde Signaling and Beyond. Annu Rev Plant Biol. 2016;67(1):25–53.

48. Zhao C, Wang Y, Chan KX, Marchant DB, Franks PJ, Randall D, et al. Evolution of chloroplast retrograde signaling facilitates green plant adaptation to land. Proc Natl Acad Sci. 2019;116(11):5015–20.

